# Deploying an *in vitro* gut model to assay the impact of a mannan-oligosaccharide prebiotic, Bio-Mos® on the Atlantic salmon (*Salmo salar*) gut microbiome

**DOI:** 10.1101/2021.08.18.456921

**Authors:** R. Kazlauskaite, B. Cheaib, J. Humble, C. Heys, U. Z. Ijaz, S. Connelly, W.T. Sloan, J. Russell, L. Martinez-Rubio, J. Sweetman, A. Kitts, P. McGinnity, P. Lyons, M.S. Llewellyn

## Abstract

Mannose-oligosaccharide (MOS) pre-biotics are widely deployed in animal agriculture as immunomodulators as well as to enhance growth and gut health. Their mode of action is thought to be mediated through their impact on host microbial communities and the associated metabolism. Bio-Mos is a commercially available pre-biotic currently used in the agri-feed industry. To assess Bio-Mos for potential use as a prebiotic growth promotor in salmonid aquaculture, we have modified an established Atlantic salmon *in vitro* gut model, SalmoSim, to evaluate its impact on the host microbial communities. Inoculated from biological triplicates of adult farmed salmon pyloric caeca compartments, the microbial communities were stabilised in SalmoSim followed by a twenty-day exposure to the prebiotic and in turn followed by an eight day ‘wash out’ period. Dietary inclusion of MOS resulted in a significant increase in formate (p=0.001), propionate (p=0.037) and isovalerate (p=0.024) levels, correlated with increased abundances of several, principally, anaerobic microbial genera (*Fusobacterium*, *Agarivorans*, *Pseudoalteromonas*). DNA metabarcoding with the 16S rDNA marker confirmed a significant shift in microbial community composition in response to MOS supplementation with observed increase in lactic acid producing *Carnobacterium*. In conjunction with previous *in vivo* studies linking enhanced volatile fatty acid production alongside MOS supplementation to host growth and performance, our data suggests that Bio-Mos may be of value in salmonid production. Furthermore, our data highlights the potential role of *in vitro* gut models to augment *in vivo* trials of microbiome modulators.

**Importance:** In this paper we report the results of the impact of prebiotic (MOS supplementation) on microbial communities within recently developed Atlantic salmon gut microbiome *in vitro* simulator. Our data suggest that Bio-Mos may be of value in salmonid production as it enhances volatile fatty acid production in the Atlantic salmon gut and correlates with a significant shift in microbial community composition with observed increase in lactic acid producing *Carnobacterium*. In conjunction with previous *in vivo* studies linking enhanced volatile fatty acid production alongside MOS supplementation to host growth and performance, our data suggest that Bio-Mos may be of value in salmonid production. Furthermore, our data highlights the potential role of *in vitro* gut models to augment *in vivo* trials of microbiome modulators.

## 1. Introduction

Since the late 1970s, the salmon aquaculture sector has grown significantly, currently exceeding 1 million tonnes of salmon produced per year (FAO, 2018). In aquaculture environments, fish are exposed to abiotic conditions and biotic interactions that are extensively different from the wild, such as changes in temperature and salinity, and close contact between animals that can favour potential disease outbreaks r (Kennedy et al., 2016), as well as long term stress through physical aggression and overcrowding (Adams et al., 2007; Turnbull et al., 2005). The rapid expansion of the aquaculture sector requires a means to promote efficient feed conversion, reduce the need for medical treatments and reduce waste discharges whilst also improving farmed fish quality.

In order to mitigate disease outbreaks and improve feed conversion, prebiotics are widely deployed in agriculture and aquaculture settings (Markowiak & Ślizewska, 2018; Patterson & Burkholder, 2003; Ringø et al., 2010). Prebiotics are defined as non-digestible food additives that have a beneficial effect on the host by stimulating growth and activity of bacterial communities within the gut that improve animal health (Gibson, Glenn R. and Roberfroid, 1995). One prebiotic type found in aquaculture is mannooligosaccharides (MOS), these are made of glucomannoprotein-complexes derived from the outer layer of yeast cell walls (*Saccharomyces cerevisiae*) (Merrifield et al., 2010). MOS compounds were shown to improve gut function and health by increasing villi height, evenness and integrity in chickens (Hooge, 2004; Iji et al., 2001), cattle (Castillo et al., 2008) and fish (A. Dimitroglou et al., 2009). MOS supplementation in monogastrics has been reported to drive changes in host associated microbial communities (Halas & Nochta, 2012; Sims et al., 2004). Associated increase of volatile fatty acid (VFA) production was reported which can have beneficial knock-on effects in terms of host metabolism and gut health (Besten et al., 2013).

There are limited number of studies investigating the effect of MOS on the fish microbiome (A. Dimitroglou et al., 2009; Ringø et al., 2016) with disparities in the observed results that could be partially explained by the duration of MOS supplementation, fish species, age or environmental conditions. For example, it was found that MOS supplemented diets improved growth and/or feed utilization in some studies (Buentello et al., 2010; Gültepe et al., 2011; Staykov et al., 2007; Torrecillas et al., 2013; Yilmaz et al., 2007), but others found that MOS supplementation did not affect fish performance or feed efficiency (Peterson et al., 2010; Pryor et al., 2003; Razeghi Mansour et al., 2012). Detailed studies are needed to investigate the effect of MOS supplements on the fish microbiome to enhance our understanding of the link between MOS and gut health. *In vitro* gut models offer the advantage of doing so in a replicated and controlled environment.

SalmoSim is a salmon gut simulation system that continuously maintains the microbial communities present in the intestine of marine phase Atlantic Salmon (*Salmo salar*) (Kazlauskaite et al., 2020). The current study deploys a modified version of SalmoSim designed to evaluate the effect of Bio-Mos (Alltech), a commercially available MOS product, on the microbial communities of the Atlantic salmon small intestine (pyloric caecum) in biological triplicate. We assayed microbial composition and fermentation in the SalmoSim system and show a significant impact of Bio-Mos supplementation on both.

## 2. Materials and Methods

### 2.1. *In vivo* sample collection and *in vitro* system inoculation

Three adult Atlantic salmon gut samples were collected from the MOWI processing plant in Fort William, Scotland and transferred to the laboratory in an anaerobic box on ice. Samples were placed in an anaerobic hood and contents from pyloric caeca compartment were scraped and collected into separate sterile tubes. Half of each sample was stored in −80°C freezer (as a backup, in case the run needed restarting), whilst the other half was used as an inoculum for the SalmoSim system. Inoculums were prepared for the *in vitro* trial from the pyloric caeca of different individual fish (three biological replicates). Prior to inoculation, inoculums were dissolved in 1 ml of autoclaved 35 g/L Instant Ocean® Sea Salt solution.

### 2.2. SalmoSim *in vitro* system preparation

*In vitro* system feed media was prepared by combining the following for a total of 2 litres: 35 g/L of Instant Ocean® Sea Salt, 10 g/L of the Fish meal (the same feed as used in (Kazlauskaite et al., 2020)), 1 g/L freeze-dried mucous collected from the pyloric caecum, 2 litres of deionised water and 0.4% of Bio-Mos (derived from the outer cell wall of *Saccharomyces cerevisiae* strain 1026) for the prebiotic supplemented feed. A supplementation level of 0.4% weight by volume was chosen based on previous studies (Arkadios Dimitroglou et al., 2011; Torrecillas et al., 2015). This feed was then autoclave-sterilised, followed by sieving of the bulky flocculate, and finally subjected to a second round of autoclaving. System architecture was prepared as described previously with some modifications (Kazlauskaite et al., 2020). In short, appropriate tubes and probes were attached to a two-litre double-jacketed bioreactor, and three 500 ml Applikon Mini Bioreactors. Four 1 cm^3^ aquarium sponge filters were added to each Mini Bioreactor vessel which were then autoclaved, sterilised, and connected as in Figure 1. Nitrogen gas was periodically bubbled through each vessel to maintain anaerobic conditions. The two-litre double jacketed bioreactor and three 500 ml bioreactors were filled with 1.5 litres and 400 ml of feed media respectively. Once the system was set up, media transfer, gas flow and acid/base addition were undertaken for twenty-four hours axenically in order to stabilise the temperature, pH, and oxygen concentration with respect to levels measured from adult salmon. SalmoSim system diagram is visualised in Figure 1. Physiochemical conditions within the three 500 ml SalmoSim gut compartments were kept similar to the values measured *in vivo* (Kazlauskaite et al., 2020): temperature inside the reactor vessels was maintained at 12°C, dissolved oxygen contents were kept at 0% by daily flushing with N_2_ gas for 20 minutes, and pH 7.0 by the addition of 0.01M NaOH and 0.01M HCl. The 2-litre double jacketed bioreactor (represents a sterile stomach compartment) was kept at 12°C and pH at 4.0 by the addition of 0.01M HCl. During this experiment (apart from the initial pre-growth period), the transfer rate of slurry between reactor vessels was 238 ml per day. Finally, on a daily basis, 1 ml of filtered salmon bile and 0.5 ml of autoclaved 5% mucous solution were added to the three bioreactors simulating pyloric caecum compartments.

**Figure 1.**
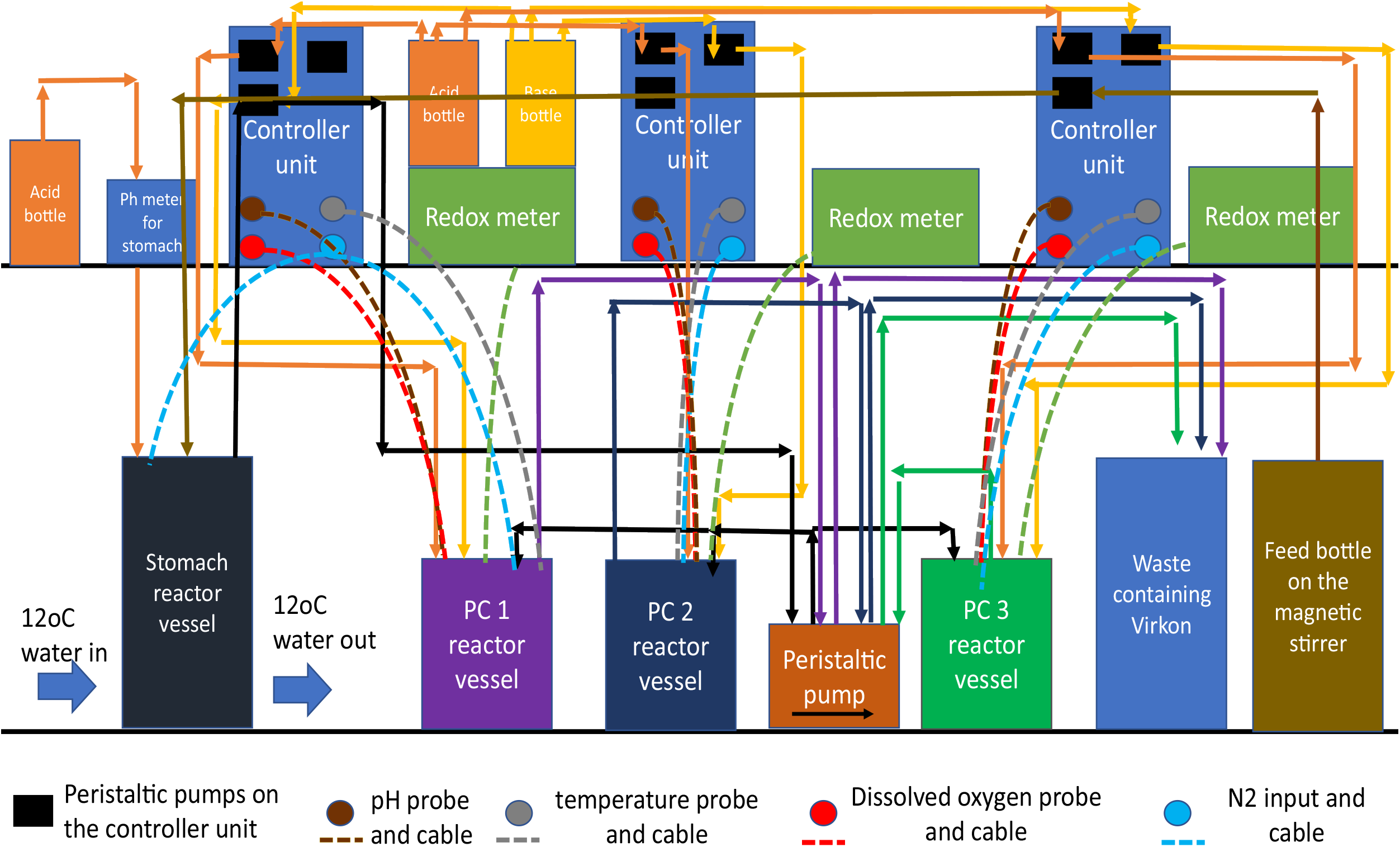
Artificial gut model system set-up and in vitro trial set up. A SalmoSim system designed to run in biological triplicate.

### 2.3. SalmoSim inoculation and microbial growth

To generate stable and representative microbial communities for experimentation (Kazlauskaite et al., 2020), microbial communities were grown within SalmoSim system during a separate twenty-four day run prior to the main experimental run (Figure 2A). This was achieved by adding fresh inoculum from pyloric caeca to three 500 ml bioreactors which was then pre-grown for 4 days without media transfer, followed by 20 days feeding the system at a 238 ml per day feed transfer rate. A volume of 15 ml of the stable communities were collected at the end of this pre-growth period, centrifuged at 3000 *g* for 10 minutes and supernatant removed. The pellet was then dissolved in 1 ml of autoclaved 35 g/L Instant Ocean® Sea Salt solution, flash frozen in liquid nitrogen for 5 minutes and stored long term in −80°C freezer.

**Figure 2.**
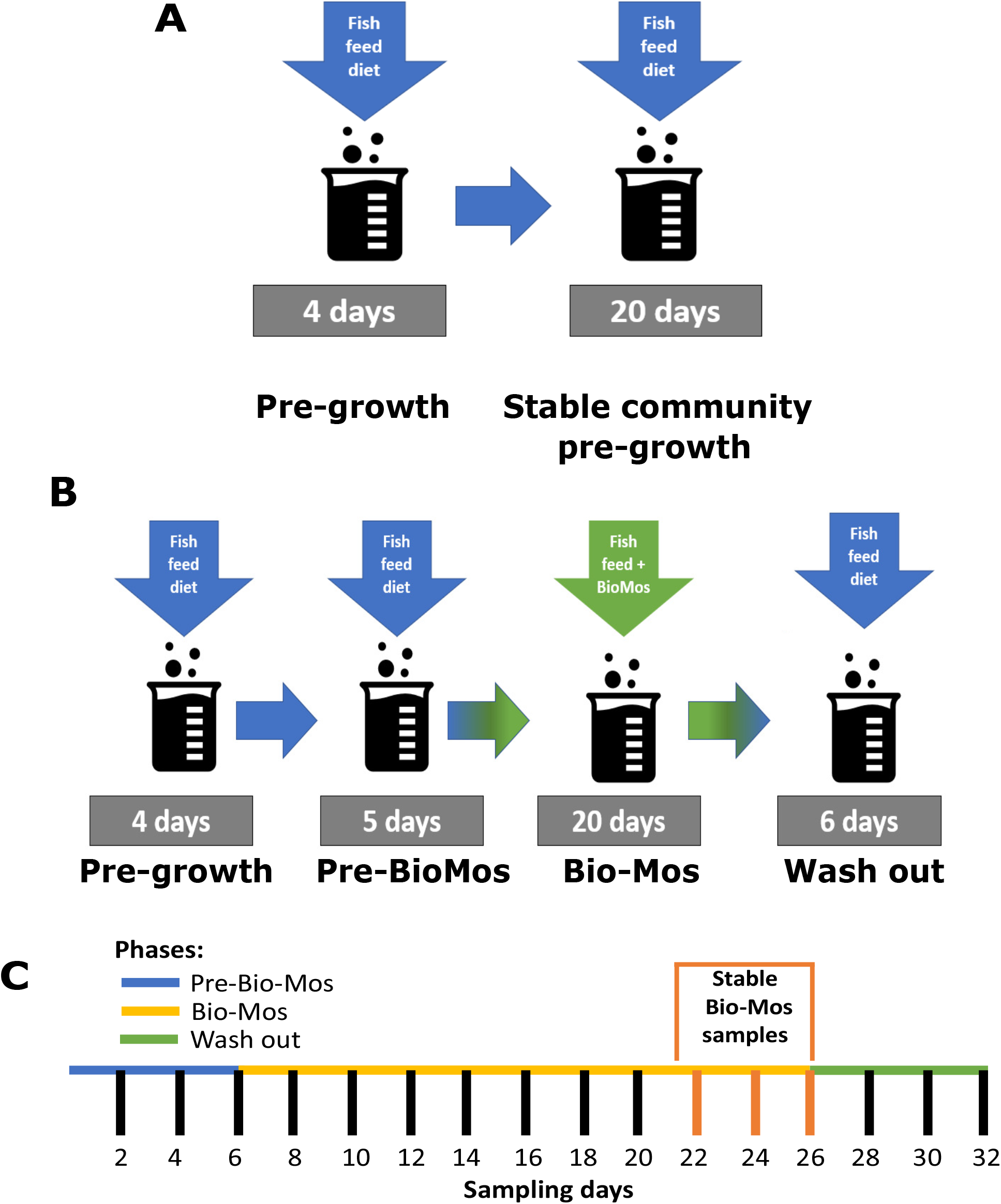
In vitro trial setup. 2A Stable community pre-growth run within the SalmoSim system; 2B Main experimental run that involved four stages: (i) pre-growth (without feed transfer for 4 days), (ii) feeding system with Fish meal (Pre-Bio-Mos: 5 days), (iii) feeding system with Fish meal diet supplemented with Bio-Mos (Bio-Mos: 20 days), (iv) wash out period during which system was fed Fish meal without the addition of prebiotic (Wash out: 6 days); 2C SalmoSim sampling time points, which include definition of stable time points for Bio-Mos phase (days 22, 24, and 26 - once bacterial communities had time to adapt to Bio-Mos addition).

### 2.4. Assaying Bio-Mos impact on microbial communities in the SalmoSim *in vitro* system

Frozen pre-grown stable pyloric caeca samples were thawed on ice and added to the SalmoSim system with each 500 ml bioreactor inoculated using bacterial communities pre-grown from a different fish. The system was run in several stages: (i) 4-day initial pre-growth period without feed transfer (Pre-growth), (ii) 5-day period during which SalmoSim was fed without prebiotic (Pre-Bio-Mos), (iii) 20-day period during which SalmoSim was fed on feed supplemented with Bio-Mos (Bio-Mos), (iv) 6-day wash out period during which SalmoSim was fed on Fish meal diet without addition of prebiotic (Wash out). The schematic representation of the experimental design is visually represented in Figure 2B. Sixteen samples were collected throughout the experimental run as described previously (Figure 2C) (Kazlauskaite et al., 2020).

### 2.5. Genomic DNA extraction and NGS library preparation

DNA extraction and NGS library preparation protocols were previously described (Heys et al., 2020; Kazlauskaite et al., 2020). Briefly, the samples collected from SalmoSim system and stable pre-grown inoculums were thawed on ice and exposed to bead-beating step for 60 seconds by combining samples with MP Biomedicals^™^ 1/4” CERAMIC SPHERE (Thermo Fisher Scientific, USA) and Lysing Matrix A Bulk (MP Biomedicals, USA). Later, DNA was extracted by using the QIAamp® DNA Stool kit (Qiagen, Valencia, CA, USA) according to the manufacturer’s protocol (Claassen et al., 2013). After, extracted DNA was amplified using primers targeting V1 bacterial rDNA 16s region under the following PCR conditions: 95°C for ten minutes, followed by 25 cycles at 95°C for 30 seconds, 55°C for 30 seconds and 72°C for 30 seconds, followed by a final elongation step of 72°C for 10 minutes. The second-round PCR, which enabled the addition of the external multiplex identifiers (barcodes), involved six cycles only and otherwise had identical reaction conditions to the first round of PCR. This was followed by the PCR product clean-up using Agencourt AMPure XP beads (Beckman Coulter, USA) according to the manufacturers’ protocol and gel-purification using the QIAquick Gel Extraction Kit (Qiagen, Valencia, CA, USA). Finally, the PCR products were pooled together at 10 nM concentration and sent for sequencing using the HiSeq 2500 sequencer.

### 2.6. NGS data analysis

NGS data analysis was undertaken as described previously (Kazlauskaite et al., 2020). In short, to determine microbial community stability within SalmoSim system over time, two alpha diversity metrics (effective microbial richness and evenness (effective Shannon)) were calculated to analyse using Rhea package (Lagkouvardos et al., 2017b) and visualised by using microbiomeSeq package based on phyloseq package (McMurdie & Holmes, 2013; Ssekagiri et al., 2017).

To provide an overall visualisation of microbial composition across all samples, Principal Coordinates Analysis (PCoA) was performed by using phyloseq package (Love et al., 2017; Ssekagiri et al., 2017) with Bray-Curtis dissimilarity measures calculated by using the vegdist() function from the vegan v2.4-2 package (Oksanen et al., 2013). Bray-Curtis distances were calculated for four different datasets: the full dataset (containing all biological replicates together), and three different subsets each containing only one of the three biological replicate samples from SalmoSim: Fish inoculum 1, 2, or 3.

To further compare microbial structure between various experimental phases, beta diversity was calculated for two different datasets: (i) all (completed data set containing all the samples sequenced) and (ii) subset (containing all samples for Pre-Bio-Mos and Wash out period, but only stable samplings from Bio-Mos period (time points 22, 12 and 26)). From these datasets ecological distances were computed using Bray-Curtis and Jaccard distances with vegdist() function from the vegan v2.4-2 package (Oksanen et al., 2013). Furthermore, the phylogenetical distances were computed for each dataset using GUniFrac() distance (generalised UniFrac) at the 0% (unweighted), 50% (balanced) and 100% (weighted) using the Rhea package (Lagkouvardos et al., 2017a). Both ecological and phylogenetical distances were then visualised in two dimensions by Multi-Dimensional Scaling (MDS) and non-metric MDS (NMDS) (Anderson, 2001). Finally, a permutational multivariate analysis of variance (PERMANOVA) was performed using distance matrices (including phylogenetic distance) to explain sources of variability in the bacterial community structure as result of changes in recorded parameters (Anderson, 2001).

To identify differentially abundant OTUs between various experimental phases (Pre-Bio-Mos, Bio-Mos and Wash out), differential abundance was calculated using microbiomeSeq package based on DESeq2 package (Love et al., 2017; Ssekagiri et al., 2017). Results were then summarised using bar plots at genus level, identifying number of OTUs belonging to specific genus level that increase or decrease between various experimental phases.

To identify OTUs that correlated with measured VFAs, the Pearson correlation coefficient (r>0.8) was calculated between taxonomic variables (OTUs) measured VFA values measured, and visualised using tools supplied by Rhea package within different experimental phases (Pre-Bio-Mos, Bio-Mos, and Wash out) (Lagkouvardos et al., 2017a).

Finally, in order to analyse microbial community structure within different experimental phases, network analysis using Spearman correlation (r>0.8) was performed on three datasets: (i) all Pre-Bio-Mos samples, (ii) stable Bio-Mos samples (samples from days 22, 24 and 26), and (iii) all Wash out samples. Key network characteristics were compared between the three experimental phases: i.e., degree and centrality betweenness. All these comparisons were analysed and visualised using “ggstatsplot” package.

### 2.7. Protein fermentation and Volatile Fatty Acid (VFA) analysis

At each sampling point, microbial protein fermentation was assessed by measuring the protein concentration using Thermo Scientific^™^ Pierce^™^ BCA Protein Assay Kit (Thermo Fisher Scientific, USA) and the ammonia concentration using Sigma-Aldrich® Ammonia Assay Kit (Sigma-Aldrich, USA). Both methods were performed according to manufacturer protocol by using a Jenway 6305 UV/Visible Spectrophotometer (Jenway, USA). For VFA analysis, nine samples from each pyloric caecum compartment were collected (from 3 biological replicates): 3 samples from Pre-Bio-Mos period (days 2-6), 3 samples from stable time points from the period while SalmoSim was fed on feed supplemented with Bio-Mos (days 22-26), and 3 samples from the Wash out period (days 28-32). VFA sampling was performed as described previously (Kazlauskaite et al., 2020). Extracted VFAs were sent for gas chromatographic analysis at the MS-Omics (Denmark).

In order to establish whether VFA concentrations were statistically different between different experimental phases (Pre-Bio-Mos, Bio-Mos and Wash out), a linear mixed effect model was deployed (Model 1) considering time point (sampling time point) and run (biological replicate of SalmoSim system) as random effects.

*Model 1* = *lmer(<u>VFA</u>~ Phase*+*(1*|*Time point)*+*(1*|*Run))*

Finally, in order to establish whether ammonia production changed throughout experimental run, a linear mixed effect model was deployed (Model 2) treating run biological replicate (of SalmoSim system) as random effect.

*Model 2* = *lmer(<u>ammonia concentration</u>~ Time point*+*(1*|*Run))*

## 3. Results

In order to explore the impact of the Bio-Mos prebiotic on microbial communities in SalmoSim, microbial amplicons in different experimental phases (Pre-Bio-Mos, Bio-Mos and wash out) were surveyed using Illumina NovaSeq amplicon sequencing of the 16S V1 rDNA locus. In total 11.5 million sequence reads were obtained after quality filtering. Alpha diversity metrics (Effective richness in Figure 3A and effective Shannon diversity in Figure 3B) indicated that the initial inoculum contained the lowest number of OTUs and had the lowest bacterial richness compared to later sampling time points from SalmoSim system, but these differences were not statistically significant. Furthermore, this figure indicates no statistically significant differences between different experimental phases (Pre-Bio-Mos, Bio-Mos and Wash out) in both terms of effective richness and Shannon diversity. Taken together, diversity and richness estimates suggest non-statistically significant increase of in the number of detectable microbial taxa as a result of transfer into SalmoSim system, but overwise stable diversity and richness over the different experimental phases.

**Figure 3.**
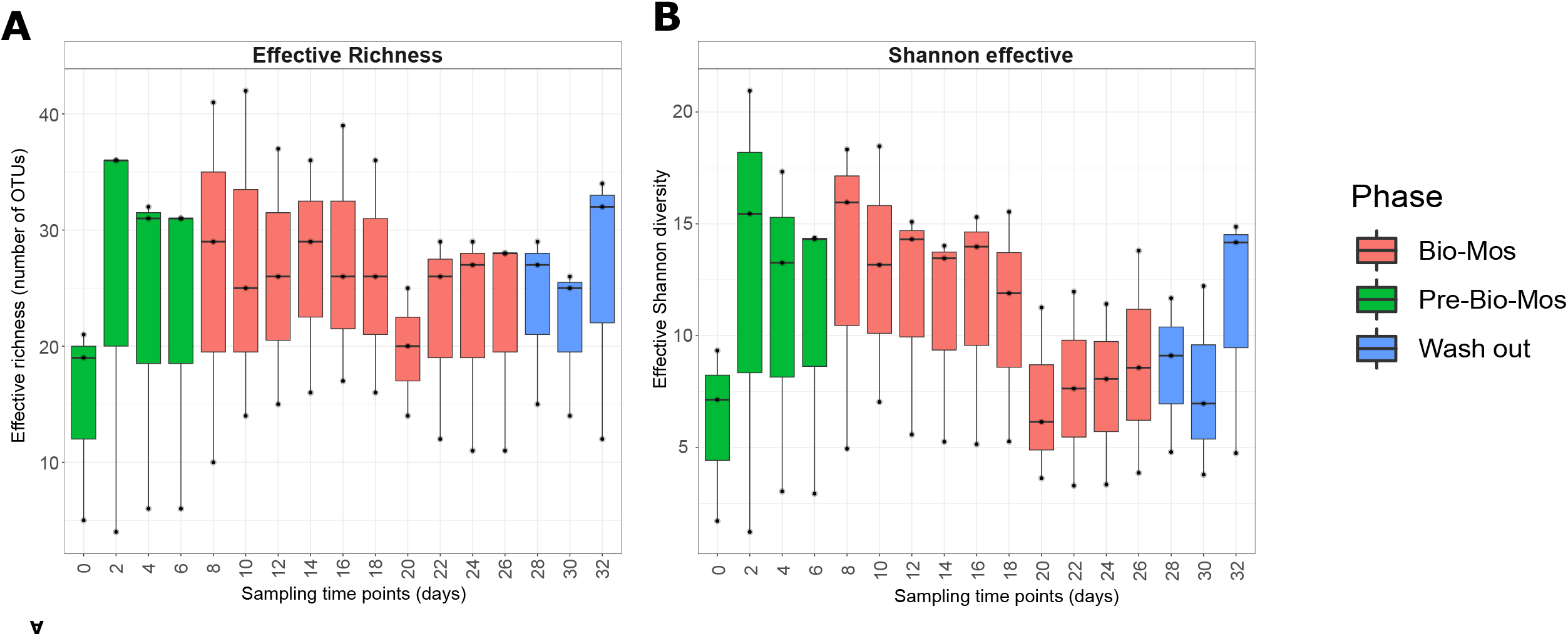
Alpha-diversity dynamics within the SalmoSim system during exposure to Bio-Mos prebiotics. The figure represents different alpha diversity outputs at different sampling time points (days) from SalmoSim system. Time point 0 represents microbial community composition within initial SalmoSim inoculum from the pre-grown stable bacterial communities, time points 2-6 identifies samples from SalmoSim system fed on Fish meal diet alone (Pre-Bio-Mos: green), time points 8-26 identifies samples from SalmoSim system fed on Fish meal diet with addition of Bio-Mos (Bio-Mos: red), and time points 28-32 identifies samples from wash out period while SalmoSim was fed on feed without addition of prebiotic (Wash out: blue). **A** visually represents effective richness (number of OTUs), **B** represents effective Shannon diversity.

To provide an overview of microbial composition and variation in the experiment, a PCoA plot was constructed based on Bray-Curtis distanced between samples (Figure 4 A-D). Biological replicate (the founding inoculum of each SalmoSim run) appears to be a major driver of community composition in the experiment (Figure 4A). This is supported by Figure 5 that visually represents varying microbial composition within different fish. Only when individual SalmoSim replicates were visualised separately in PCoA plots, do the changes to microbial communities in response to the different experimental phases become apparent (Figures 4 B-D). These results indicate that bacterial communities shift from Pre-Bio-Mos to Bio-Mos, but they remain fairly stable (statistically similar, p>0.05 in majority of cases) between Bio-Mos and Wash out periods as reflected by beta diversity results summarised in Supplementary Table 1. However, community shifts do not necessarily occur along the same axes in each SalmoSim replicate indicative, perhaps, or a different microbiological basis for that change. This trend is confirmed in Figure 5 that indicates a more substantial shift in microbial community profile between Pre-Bio-Mos and Bio-Mos phases in Fish 2 and 3, but to the lesser extent in Fish 1. Results were further confirmed by performing beta-diversity analysis using both phylogenetic and ecological distances, both of which indicated statistically significant differences between Pre-Bio-Mos and Bio-Mos phases, but not between Bio-Mos and Wash out periods (Supplementary Table 1). Furthermore, Supplementary Table 1 indicates that 149 OTUs were found to be differentially abundant between Pre-Bio-Mos and Bio-Mos phases, while only 5 OTUs were differentially abundant between Bio-Mos and Wash out phases.

**Figure 4.**
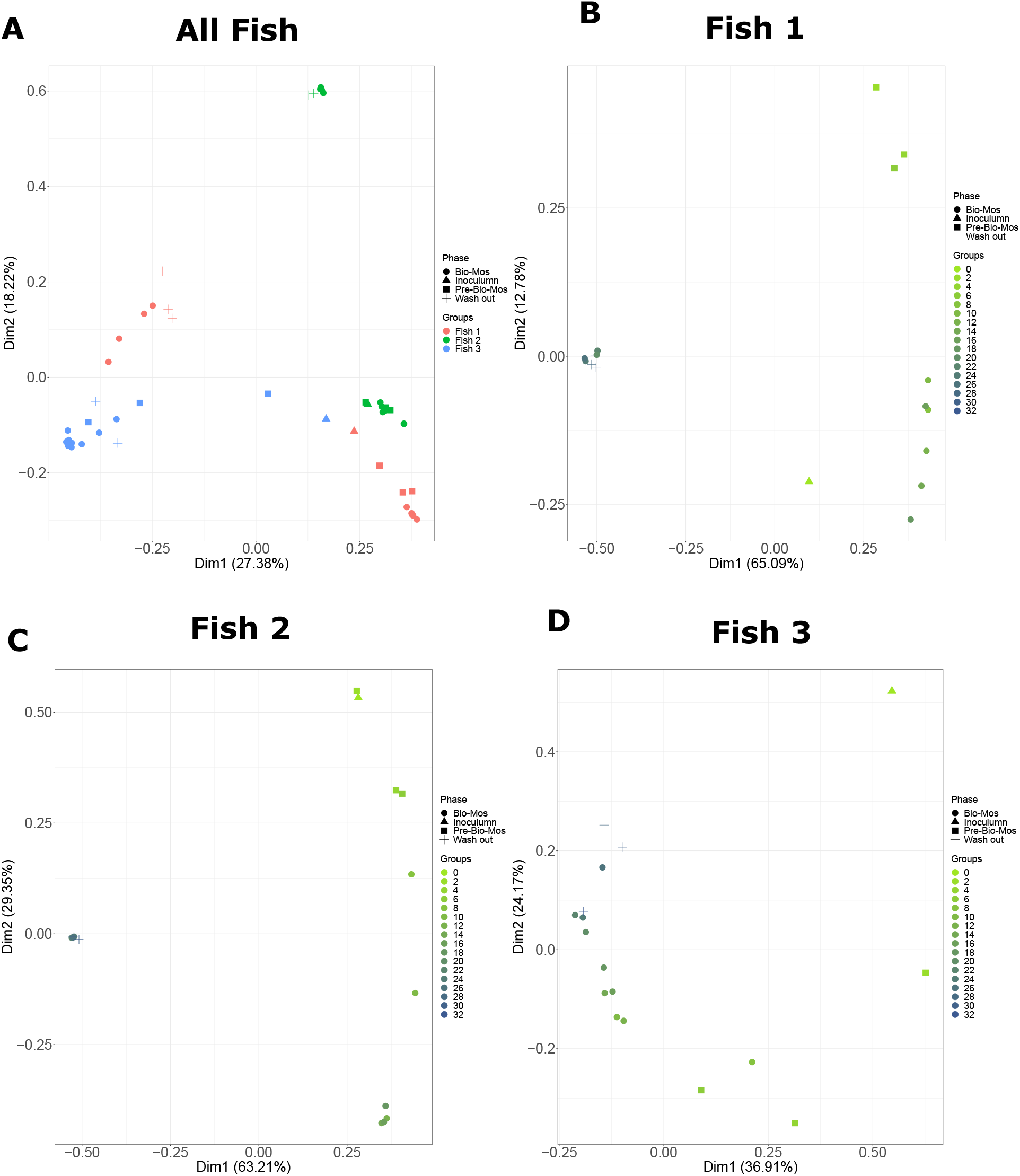
Beta diversity plots visualising bacterial communities dissimilarities within the SalmoSim bioreactors during exposure to Bio-Mos prebiotic. In the PCoA plots, Bray-Curtis distance was used between samples originating from different experimental phases (Inoculum, Pre-Bio-Mos, Bio-Mos and Wash out), annotated with sampling time points and biological replicates. **A** represents all sequenced data together for all 3 biological replicates in which different colours represent different biological replicates (samples from pyloric caecum from 3 different fish) and different shapes represent different experimental phases (Inoculum, Pre-Bio-Mos, Bio-Mos and Wash out); **B-D** represent sequenced data for each individual biological replicate (**B**: Fish 1, **C**: Fish 2, **D**: Fish 3). In figures B-D different colours represent different sampling time points and different shapes represent different experimental phases (Inoculum, Pre-Bio-Mos, Bio-Mos and Wash out). Dim 1 is principal coordinate 1 and Dim 2 is principle coordinate 2.

**Figure 5.**
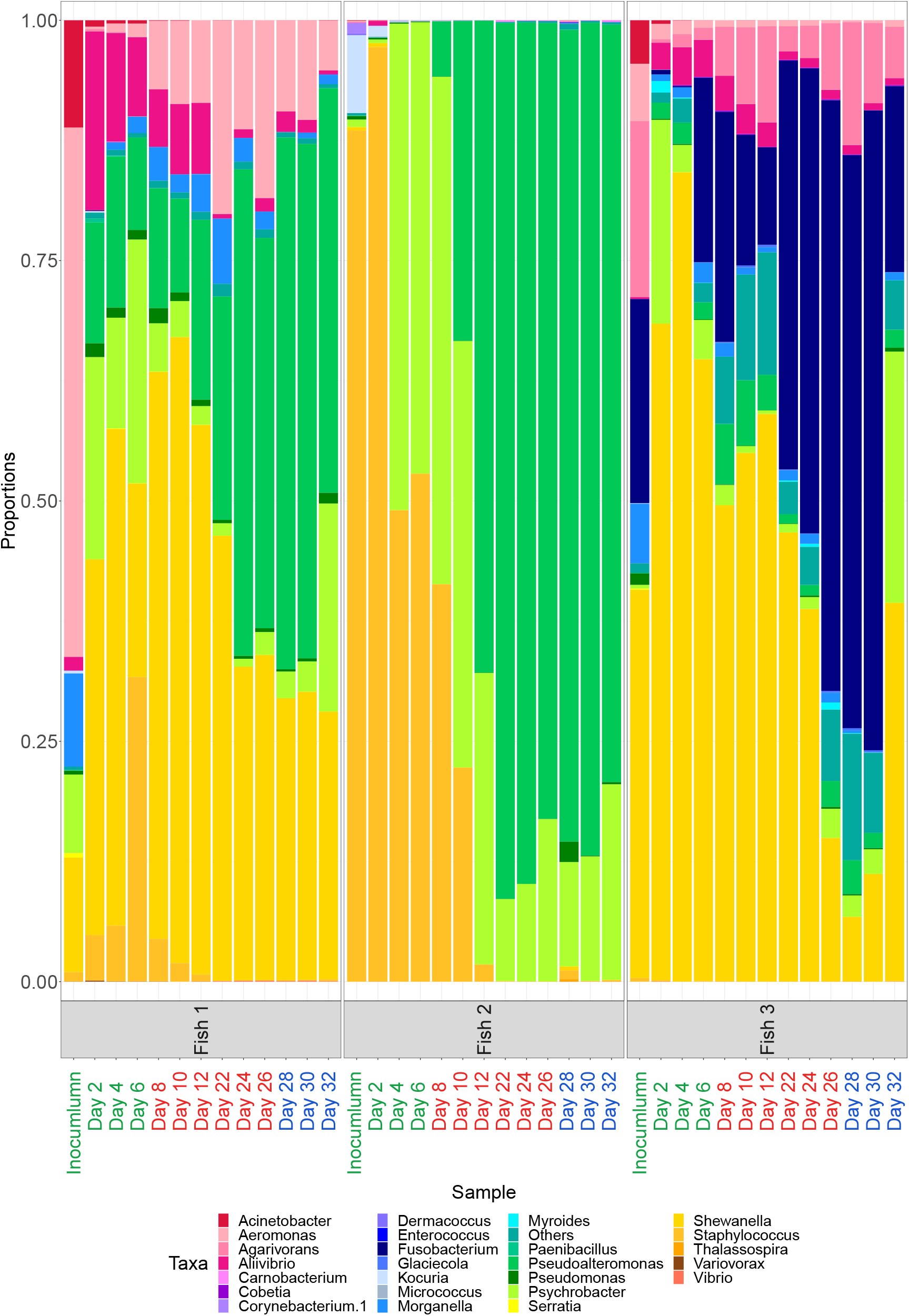
Microbial composition (25 most common genus + others) amongst different biological replicates and experimental phases. Labels on X axis in green represent samples from Pre-Bio-Mos phases, in red samples fed on Bio-Mos phase and in blue samples from Wash out period. Only subset of time points is visualised for each phase: time points 2-6 for Pre-Bio-Mos, 8-12 and 22-24 for Bio-Mos, and 28-32 for Wash out.

To compare experimental phases in more detail, differentially abundant OTUs between various experimental phases were summarised in bar plots at genus level in Figure 6. The Figure 6 A indicates that between Pre-Bio-Mos and Bio-Mos phases, more OTUs decreased in abundance, rather than increased. The OTUs that differentially increased from Pre-Bio-Mos to Bio-Mos phase were identified to belong to: *Aeromonas* (higher proportion increased (50%) rather than decreased (12.5%)), *Agarivorans*, *Aliivibrio*, *Carnobacterium* (only showed increase and no decrease), *Fusobacterium*, *Pseudoalteromonas*, *Pseudomonas*, *Psychobacter*, and *Shewanella*. Figure 6 B indicates the increase of OTUs belonging to *Enterococcus* and *Thalassospira* genera between Bio-Mos and Wash out, while OTUs belonging to *Micrococcus*, *Myroides* and *Shewanella* genera have decreased.

**Figure 6.**
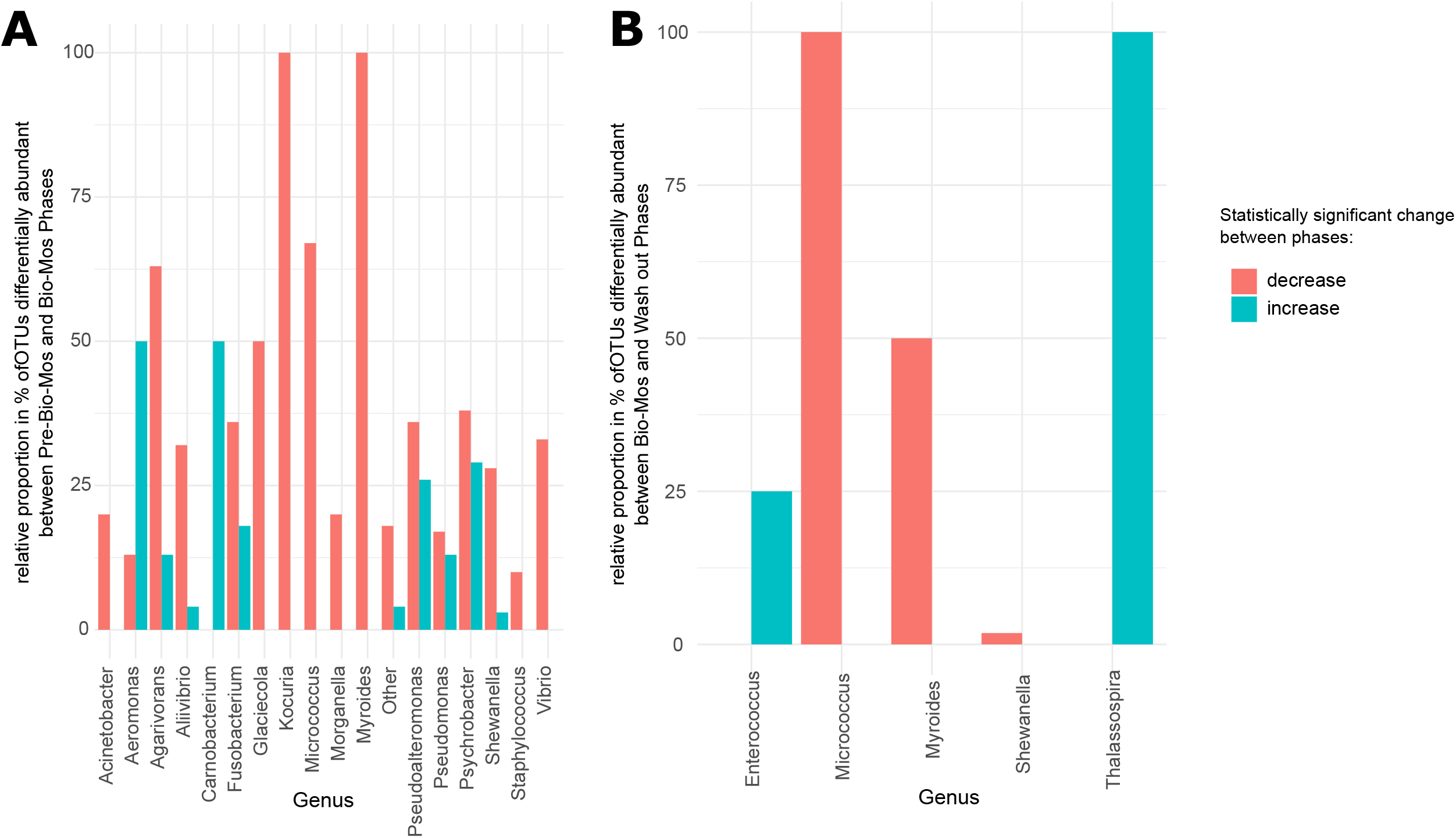
Differential abundance of OTUs grouped at genus level between different experimental phases (Pre-Bio-Mos, Bio-Mos and Wash out) Differential abundant OTUs grouped at genus level between different experimental phases: Pre-Bio-Mos vs Bio-Mos (A), Bio-Mos vs Wash out (B). Red and blue represents statistically significant (p<0.05) decrease and increase respectively between the experimental phases compared.

For the analysis of the microbial community structure throughout the experiment, three OTU co-occurrence networks were analysed for each phase (Pre-Bio-Mos, Bio-Mos, and Wash out), and the main network characteristics were compared: the degree and centrality betweenness (Supplementary Figure 7). Pre-Bio-Mos phase (Supplementary Figure 7A) indicates a higher average degree (number of edges per node) than in Bio-Mos or Wash out phases. However, the median of degrees is much higher in Bio-Mos phase compared to Pre-Bio-Mos, suggesting that during Pre-Bio-Mos phase there were clusters of interacting OTUs (one cluster with a high degree and another with lower degree). As such, the distribution of connectivity is more uniform in Bio-Mos phases, compared to Pre-Bio-Mos. Moreover, the average of betweenness centralities (centrality measure based on the shortest paths between nodes) are higher in Bio-Mos and Wash out phases compared to Pre-Bio-Mos phase (Supplementary Figure 7B).

VFA levels were measured throughout the SalmoSim trial for the stable time points (time points 2, 6 and 8 for Pre-Bio-Mos, time points 22, 24 and 26 for Bio-Mos, and time points 28, 30 and 32 for Wash out period). These results are visually represented in Figure 7, which indicates that statistically significant increases were found between Pre-Bio-Mos and Bio-Mos phases in formic, propanoic and 3-methylbutanoic acid concentrations. No significant differences in any VFA production by the system was noted between Bio-Mos and Wash out periods.

**Figure 7.**
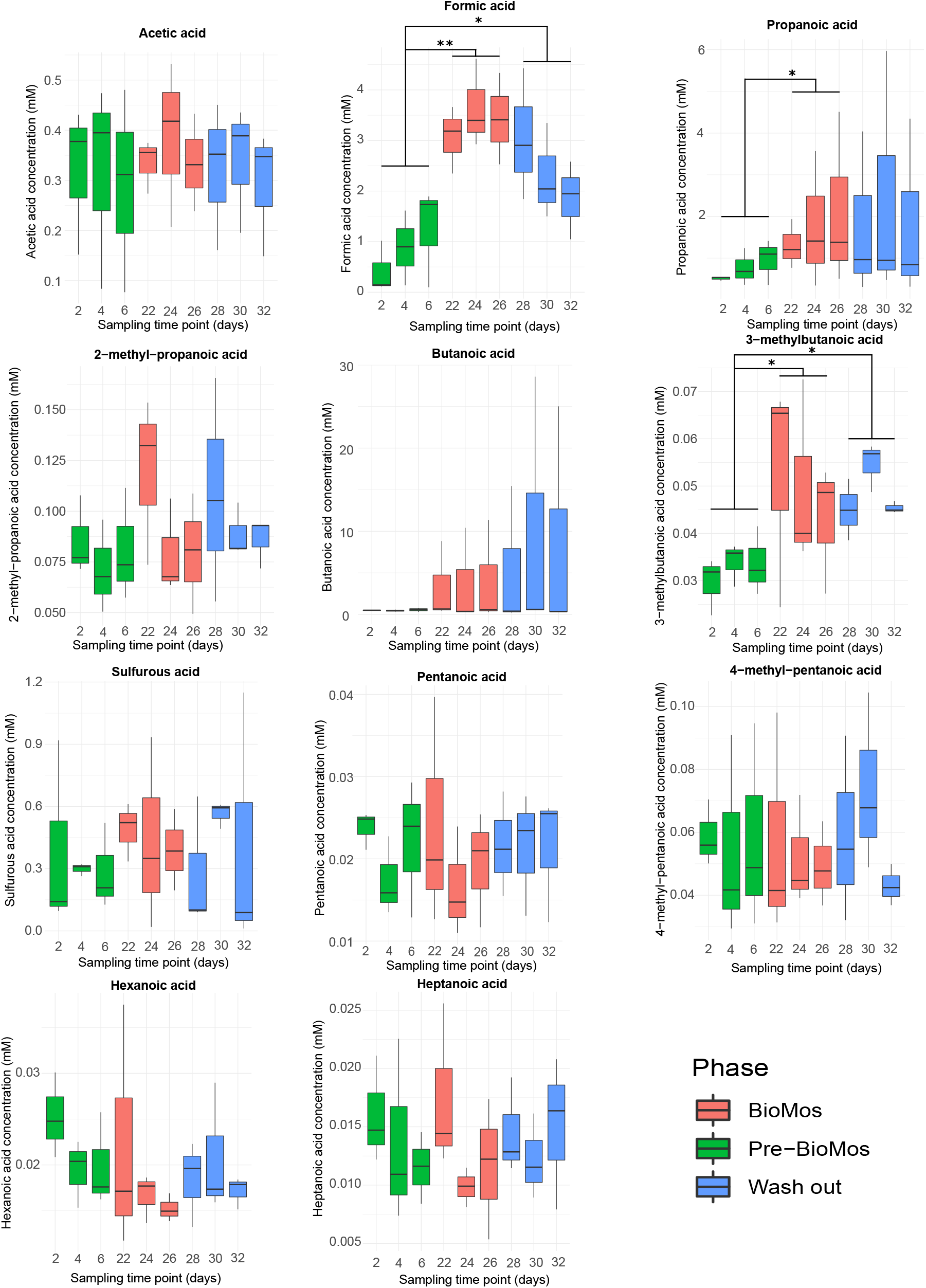
VFA responses in SalmoSim pyloric caecum compartment after Bio-Mos introduction and subsequent wash out period. The figure above visually represents 11 volatile fatty acid production in three different experimental phases: (i) SalmoSim fed on Fish meal alone without prebiotic addition (Pre-Bio-Mos: green), (ii) SalmoSim fed on Fish meal with addition of Bio-Mos (Bio-Mos: red), (iii) wash out period during which SalmoSim was fed on Fish meal without Bio-Mos (Wash out: blue). X axis represents the concentration of specific volatile fatty acid (mM) while the Y axis represents different sampling time points (days). The lines above bar plots represent statistically significant differences between different experimental phases. The asterisks show significance: (\**: 0.01* ≤ *p* < *0.05;* \*\**: 0.05* ≤ *p* < *0.001;* \*\*\**: p* ≤ *0.001)*.

Bacterial correlates of VFA increases between phases (Figure 8) were established via Pearson correlation (r>0.8). Results shown in Figure 11 identify that in the Bio-Mos phase alone, a number of OTUs which showed a strong correlation with various VFAs, had already been picked up by differential abundance analysis (Figure 6), identifying statistically significant increases. OTUs belonging to *Agarivorans* and *Fusobacterium* genera were found to be positively correlated with propanoic and formic acid, but negatively correlated with 3-methyl butanoic acid. OTU belonging to *Pseudoaltermonas* genus was found to be positively correlated with propanoic acid, but negatively correlated with 3-methyl butanoic acid, while other OTUs belonging to *Pseudoaltermonas* genus were found to be negatively correlated to propanoic acid. Finally, one OTU belonging to *Fusobacterium* was found to be negatively correlated to 3-methyl butanoic acid. Within Pre-Bio-Mos and Wash out phases, statistically significant Pearson correlations (r>0.8) were also identified between various OTUs and VFAs, however, these OTUs were not picked up in differential abundance analysis between those phases (Figure 6).

**Figure 8.**
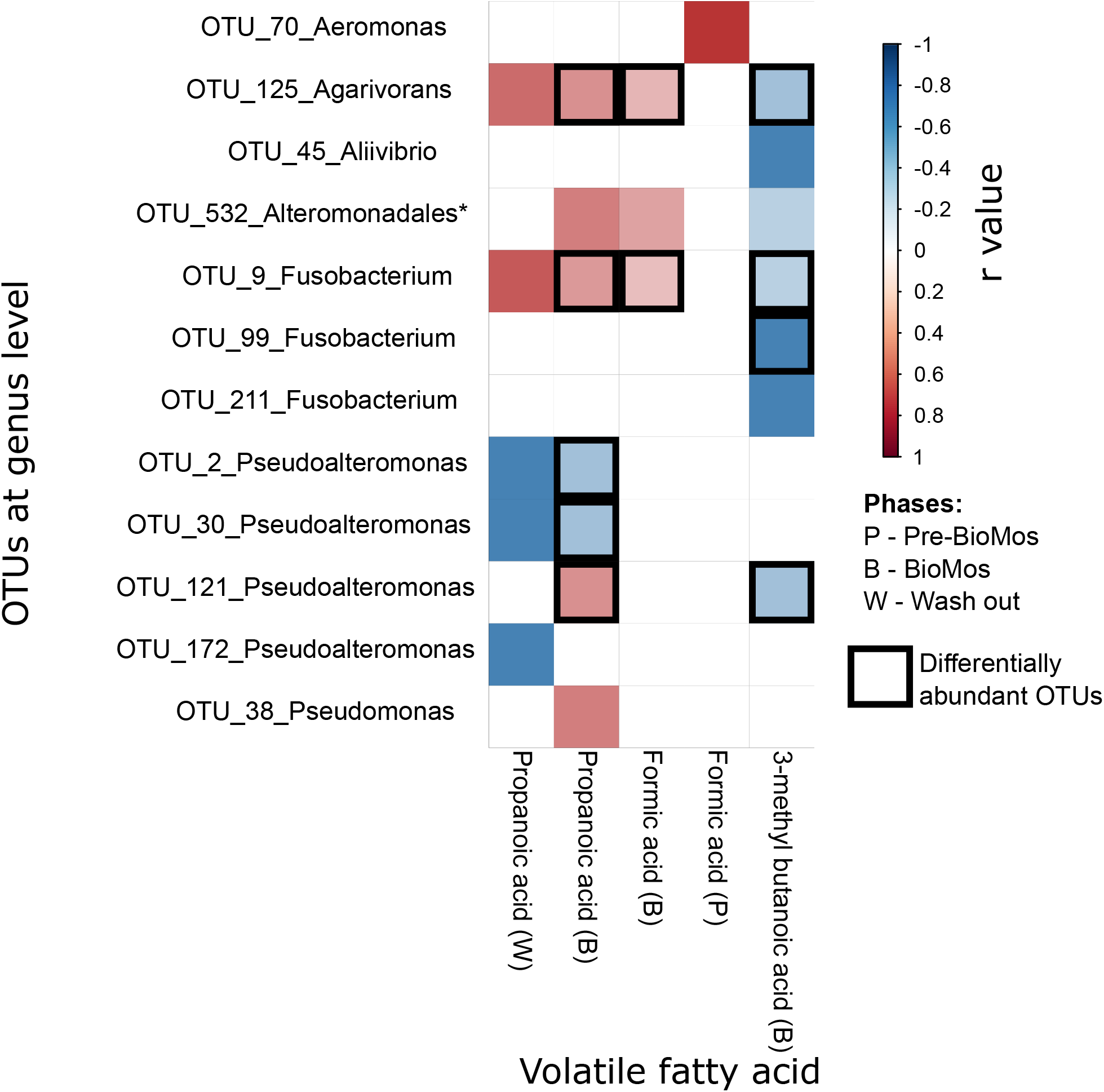
Pearson correlation coefficients across VFAs and taxonomic variables. Statistically significant (p<0.05) and strongly correlated (r>0.8) Pearson correlation coefficients across a set of VFAs (that showed statistically significant change between feeds: propanoic, formic and 3-methyl butanoic acids) and taxonomic variables (OTUs summarised at genus level apart from * to order level) are shown in various experimental phases (Pre-Bio-Mos, Bio-Mos and Wash out). Blue colour represents negative correlation and red colour represents positive correlations, respectively. The boxes indicate that these OTUs in differential abundance analysis showed statistically significant increase from Pre-Bio-Mos to Bio-Mos phase.

Figure 9 visually summarises measured ammonia (NH_3_) concentration changes through the experiment. The data indicates statistically significant increase in ammonia production between time points 2 and 4, and between time points 20 and 22, and statistically significant decrease in ammonia concentration between time points 30 and 32.

**Figure 9.**
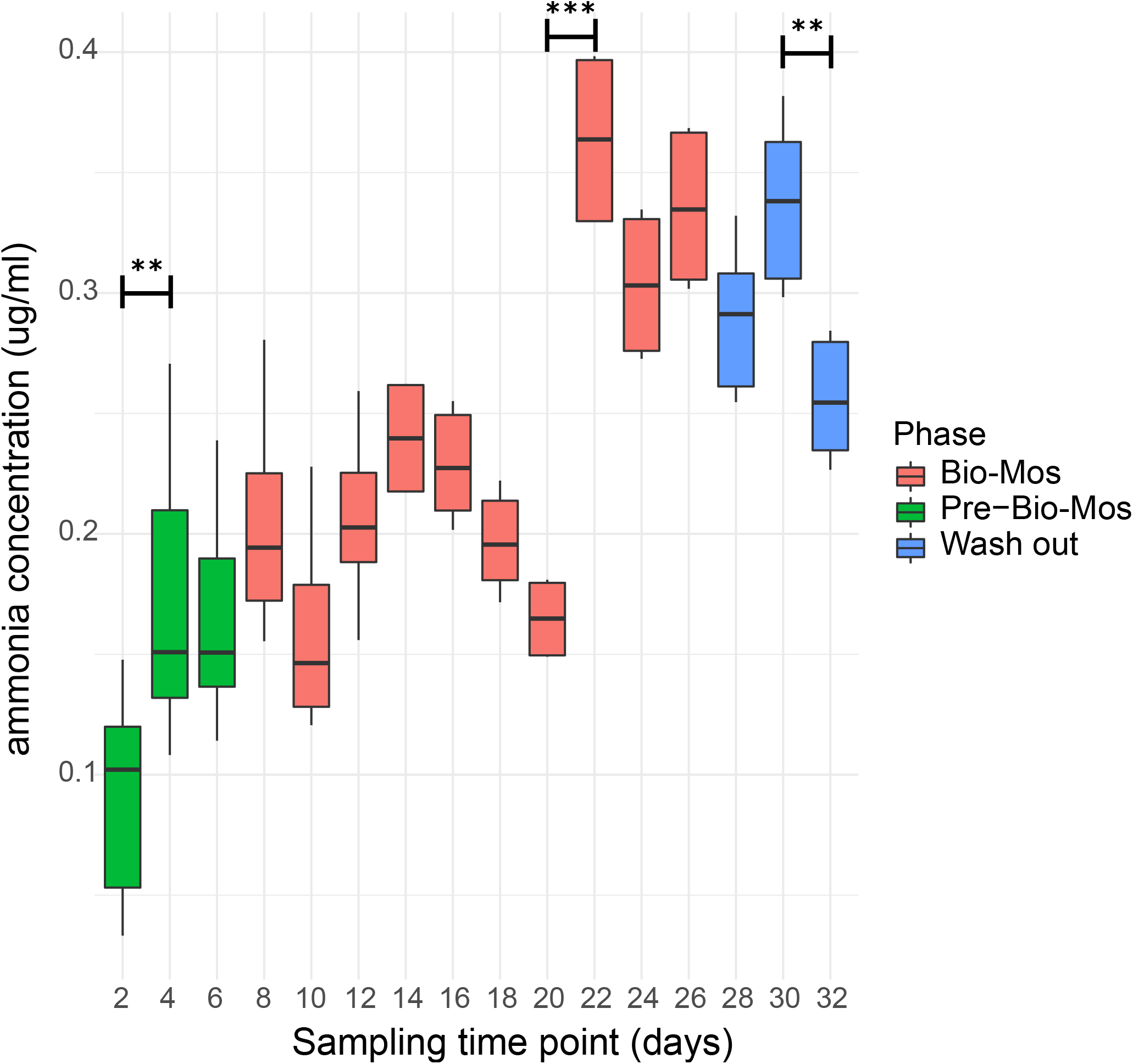
ammonia (NH_3_) concentration in SalmoSim pyloric caecum compartment throughout experiment. Ammonia (NH_3_) production in three different experimental phases: (i) SalmoSim fed on Fish meal alone without prebiotic addition (Pre-Bio-Mos: green), (ii) SalmoSim fed on Fish meal with addition of Bio-Mos (Bio-Mos: red), (iii) wash out period during which SalmoSim was fed on Fish meal without Bio-Mos (Wash out: blue). X axis represents the concentration of ammonia (μg/ml) while the Y axis represents different sampling time points (days). The lines above bar plots represent statistically significant differences between sequential time points. The asterisks show significance: (*: 0.01 ≤ p < 0.05; **: 0.05 ≤ p < 0.001; ***: p ≤ 0.001)

## 4. Discussion

Our study aimed at elucidating the effect of a commercially available MOS product (Bio-Mos) on the microbial communities within the gut content of Atlantic salmon using a newly developed artificial salmon gut simulator ‘SalmoSim’. Inclusion of Bio-Mos within the tested feed did not affect microbial community diversity and richness in the SalmoSim system, nor did subsequent removal of the prebiotic during wash out. The biological replicate (the founding inoculum of each SalmoSim run) appears to be a major driver of variations in community composition and structure throughout the experiment. This could be partially explained by the fact that feed used in the *in vitro* study was sterile, thus the bacterial communities retrieved within the SalmoSim system originated only from real salmon inoculums as in a previous experiment involving SalmoSim (Kazlauskaite et al., 2020). Our results indicate that bacterial community composition between Pre-Bio-Mos and Bio-Mos experimental phases was significantly different but was statistically similar between Bio-Mos and Wash out periods. Similar trends were observed in the bacterial activity (VFA production) that showed statistically significant increases in formic, propanoic and 3-methylbutanoic acid concentrations during the shift from Pre-Bio-Mos to Bio-Mos phase, but no statistically significant change in bacterial activity between Bio-Mos and wash-out periods. The lack of change in bacterial composition and activity between Bio-Mos and Wash out period could be explained by the short time frame of the Wash out period, lasting only 6 days, compared to the 20-day Bio-Mos phase. This is potentially not long enough to see a reversal any of changes driven by Bio-Mos. Finally, a statistically significant increase in the ammonia production during Bio-Mos phase was observed at the later time points (between days 20 and 22), followed in the reduction in ammonia concentration during Wash out period (between days 30 and 32), the potential drivers of which are discussed.

Several studies have shown that in vertebrates (e.g. chicken, mouse, turkey) supplementing feed with MOS increases the production of propionate and butyrate by gut bacteria (Ao & Choct, 2013; Pan et al., 2009; Zdunczyk et al., 2005), while other studies have not reported any effect of MOS on the VFA production (Gürbüz et al., 2010). In our study we report a statistically significant increase in the production of formic, propanoic and 3-methylbutanoic acids in the SalmoSim system associated with feed supplemented with Bio-Mos. In humans propionate is commonly absorbed and metabolised by the liver, where it impacts host physiology via regulation of energy metabolism (El Hage et al., 2020). It has also been associated with healthy gut histological development and enhanced growth in fish and shellfish (da Silva et al., 2016; Wassef et al., 2020). Formic acid, although frequently deployed as an acidifier in monogastrics to limit the growth of enteric pathogens (Luise et al., 2020), is not known to directly impact host phenotype. Similarly, except as the rare genetic disorder that occurs in humans, isovaleric acidemia, where the compound accumulates at high levels in the absence of isovaleric acid-CoA dehydrogenase activity in host tissues (Vockley & Ensenauer, 2006), isovaleric (3-methylbutanoic) acid is not expected to directly impact host phenotype either.

Further analysis identified that an increase in formic acid during the Bio-Mos phase positively correlated with OTUs belonging to *Agarivorans* (facultative anaerobic) and *Fusobacterium* (anaerobic) genera. While an increase in propanoic acid during the Bio-Mos phase also positively correlated with OTUs belonging to *Agarivorans* (facultative anaerobic) and *Fusobacterium* (anaerobic) genera as well as the *Pseudoaltermonas* (facultative anaerobic) genus, a negative correlation was found with two OTUs belonging to the same *Pseudoaltermonas* (facultative anaerobic) genus. Finally, only negative correlations were identified between the increased amount of 3-methyl butanoic acid in the Bio-Mos phase and OTUs belonging to *Pseudoaltermonas* (facultative anaerobic), *Fusobacterium* (anaerobic) and *Agarivorans* (facultative anaerobic) genera. All of these OTUs were found to not only be correlated with increased VFAs, but also to be differentially abundant between Pre-Bio-Mos and Bio-Mos phases, providing circumstantial evidence for a link between these microbes and the measured metabolites. The causal directionality between these genera and the respective VFAs is hard to establish. A strong positive correlation has been found previously in humans between the *Fusobacterium* genus and propanoic acid concentration (Riordan, 2007). Propionate is a substrate that can be metabolised by several classes of methanogenic anaerobes (Mah et al., 1990) and may be driving the growth of the genera noted here. Equally, propionate is a major product of microbial metabolism of amino acids (Louis & Flint, 2017), and it is likely here that more efficient protein metabolism in the system by certain genera is driving its abundance. An increase in ammoniacal nitrogen (ammonia) production was noted after the addition of Bio-Mos in all three replicates, albeit with a noticeable lag. Furthermore, although formate, propionate, isovalerate and ammonia show a downward trend after the removal of Bio-Mos, seemingly a longer wash-out period is required to allow VFA and ammonia to recover their pre-Bio-Mos levels.

Previously published research has suggested that feed supplementation with MOS modulates immune response in animals by stimulation of the production of mannose-binding proteins which are involved in phagocytosis and activation of the complement system (Franklin et al., 2005; Taschuk & Griebel, 2012). Such relationships with host immunity are difficult to predict with a simplified *in vitro* system. It is thought the feed supplementation with MOS elevates the immune response within the host by increasing the lactic acid bacteria (LAB) levels in common carp (Momeni-Moghaddam et al., 2015). In the present study, an increase in differential abundance of *Carnobacterium* (LAB bacteria) from Pre-Bio-Mos to Bio-Mos phases was observed. This bacterial genus has been proposed as a potential probiotic when present within Atlantic salmon (*Salmo salar*) and rainbow trout (*Oncorhynchus mykiss*) (Robertson et al., 2000). The use of Carnobacteria as probiotics were shown to be correlated with increased survival of the larvae of cod fry and Atlantic salmon fry (Gildberg et al., 1995), rainbow trout (Irianto & Austin, 2002), and salmon (Robertson et al., 2000). A fishmeal-based diet with limited carbohydrate content was used to perform this experiment, and has been previously linked to lower abundances of lactic acid producing bacteria when compared to microbial gut composition of Atlantic salmon fed on plant-based feed (Reveco et al., 2014). To enhance LAB growth even further alongside MOS in protein rich diets, some carbohydrate supplementation may be necessary.

Network analysis suggested a change in the distribution of connectivity of the microbial network during the Bio-Mos phase as compared to the Pre-Bio-Mos phase. The microbial network during the Bio-Mos phase shows higher modularity (nodes in the network tend to form denser modules), that is also reflected by a higher average of betweenness centralities within the Bio-Mos phase, a measure which represents the degree of interactive connectivity between nodes. Thus, feed supplementation with Bio-Mos may be correlated with more frequent species-species interactions, and a greater stability of network structure within the network. Stable microbial communities are also though to contribute to pathogen colonisation resistance via nutrient niche occupancy (Romero et al., 2014; Stecher et al., 2013; Xiong et al., 2019). However, a challenge experiment would be required to test this assertion.

## 5. Conclusions

Our study indicates the positive correlation between Bio-Mos supplementation and production of propanoic and formic acids, both of which are known to benefit animal microbiome and health (EFSA, 2014; Haque et al., 1970). Although, our *in vitro* model lacks a host component, previous studies involving the use of gut simulators to analyse the effectiveness of various pre-biotics were shown to produce similar results to *in vivo* trials (Duysburgh et al., 2020; Sivieri et al., 2014). Our data highlights the potential usefulness of various *in vitro* gut systems in fin fish aquaculture to study the effectiveness of feed additives.

## 6. Ethics approval and consent to participate

Animals sampled in the study were euthanised by authorised MOWI employees under Home Officer Schedule 1 of the Animals (Scientific Procedures) Act 1986.

## 7. Consent for publication

Not applicable

## 8. Availability of data and material

Sequence data have been deposited alongside metadata to the NCBI Short Read Archive

## 11. Authors’ contributions

RK and ML conceived the experiment, and RK, JH, CH, JR and AK performed the *in vitro* experimental procedure and sampling. RK performed the DNA extraction and molecular biology experiments including libraries preparation and quantification. RK prepared samples for VFA analysis and analysed the results. RK and BC produced and analysed the NGS results and performed functional diversity analysis. RK and ML wrote the manuscript. All authors reviewed, edited and approved the final draft of the manuscript.

## 12. Acknowledgements

We thank Llewellyn Environmental biotechnology laboratories teams for their help in sampling. Big thanks to MOWI team in Fort William Scotland processing plant for letting us collect our samples.

---

**Supplementary Table 1.** Beta diversity and differential abundance values from the comparison of microbial composition between different phases (Pre-Bio-Mos, Bio-Mos and Wash out) The table summarises different beta-diversity analysis outputs calculated by using different distances: phylogenetic (unweighted, balanced and weighted UniFrac) and ecological (Bray-Curtis and Jaccard’s), between different experimental phases: Pre-Bio-Mos, Bio-Mos and Wash out. Numbers represent p-values, with p-values <0.05 identifying statistically significant differences between compared groups. The comparisons are shown for 3 different datasets: (i) All (completed data set containing all the samples sequenced), (ii) a Subset (containing all samples for Pre-Bio-Mos and (iii) the Wash out period, but only stable samplings from Bio-Mos period (time points 22, 24 and 26)). The last row indicates the number of differentially abundant OTUs between Phases of interest.

| Test |  | Data | Pre-Bio-Mos vs<br>Bio-Mos | Bio-Mos vs Wash<br>out | Pre-Bio-Mos vs<br>Wash out |
| --- | --- | --- | --- | --- | --- |
| UniFrac | Unweighted (0%) | All | 0.042 | 0.029 | 0.001 |
|  |  | Subset | 0.002 | 0.184 | 0.001 |
|  | Balanced (50%) | All | 0.023 | 0.207 | 0.001 |
|  |  | Subset | 0.007 | 0.648 | 0.001 |
|  | Weighted (100%) | All | 0.022 | 0.37 | 0.002 |
|  |  | Subset | 0.002 | 0.717 | 0.001 |
| Bray-Curtis |  | All | 0.034 | 0.04 | 0.001 |
|  |  | Subset | 0.003 | 0.727 | 0.001 |
| Jaccards |  | All | 0.018 | 0.05 | 0.001 |
|  |  | Subset | 0.001 | 0.8 | 0.001 |
| Differential abundance |  | Subset | 149 | 5 | 138 |

**Supplementary Figure 1.**
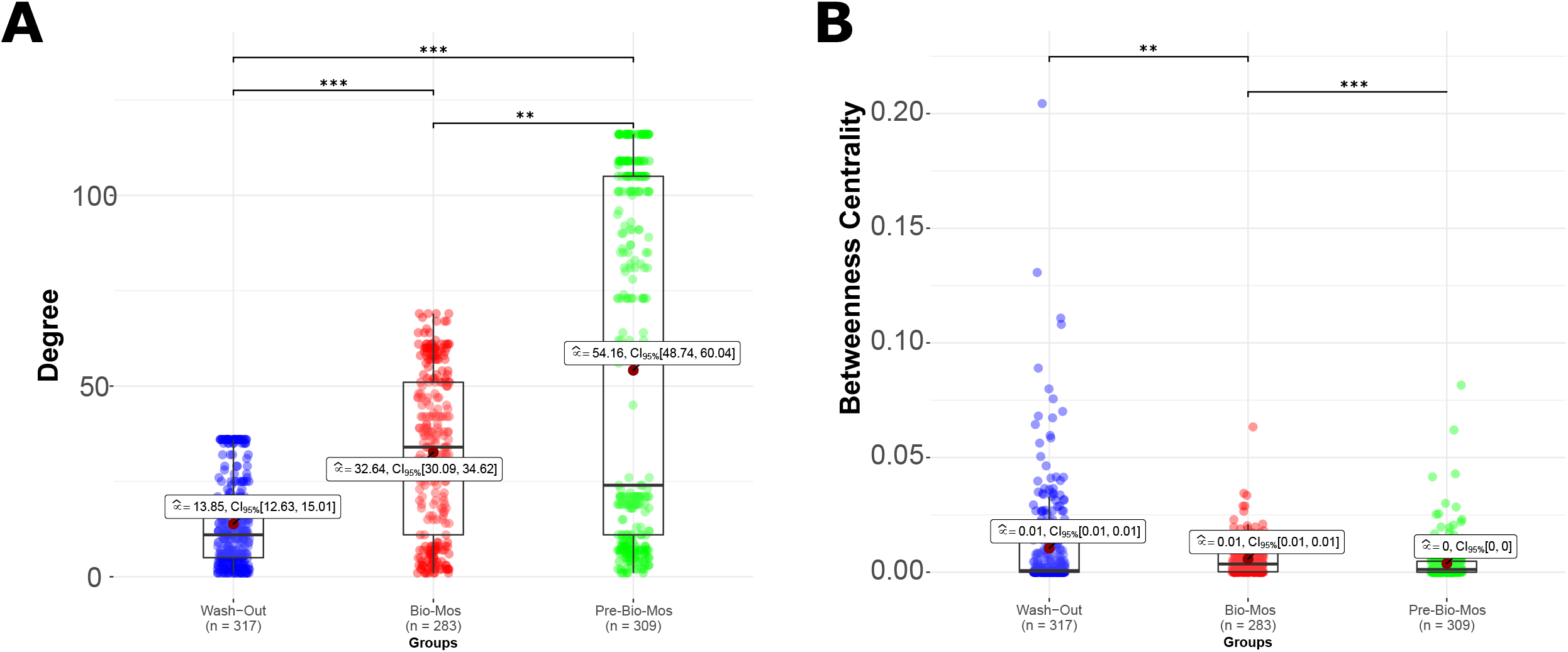
Comparison of key network analysis indicators between different experimental phases (Pre-Bio-Mos, Bio-Mos and Wash out) Figure compares key characteristics of networks produced for three experimental phases: Pre-Bio-Mos (green), Bio-Mos (red), and Wash out (blue). **A** compares degree of each network; **B** betweenness centrality. The asterisk show significance: (*: 0.01 ≤ p < 0.05; **: 0.05 ≤ p < 0.001; ***: p ≤ 0.001)

